# Molecular dissection of the chromosome partitioning protein RocS and regulation by phosphorylation

**DOI:** 10.1101/2024.07.25.605149

**Authors:** Margaux Demuysere, Adrien Ducret, Christophe Grangeasse

## Abstract

Chromosome segregation in bacteria is a critical process ensuring that each daughter cell receives an accurate copy of the genetic material during cell division. Active segregation factors such as the ParABS system or SMC complexes, are usually essential for this process but are surprisingly dispensable in *Streptococcus pneumoniae*. Rather, chromosome segregation in *S. pneumoniae* relies on the protein RocS, although the molecular mechanisms involved remain elusive. By combining genetics, *in vivo* imaging and biochemical approaches, we dissected the molecular features of RocS involved in chromosome segregation. Specifically, we investigated the respective function of the three RocS domains, specifically the C-terminal amphipathic helix (AH), the N-terminal DNA-binding domain (DBD) and the coiled-coil domain (CCD) separating the AH and the DBD. Notably, we found that a single AH is not sufficient for membrane binding and that RocS requires prior oligomerization to interact with the membrane. We further demonstrated that this self-interaction was driven by the N-terminal part of the CCD. On the other hand, we revealed that the C-terminal part of the CCD corresponds to a domain of unknown function (DUF 536) defined by three conserved glutamines which play a crucial role in RocS-mediated chromosome segregation. Finally, we showed that the DBD is phosphorylated by the unique serine-threonine kinase of *S. pneumoniae* StkP, and that mimicking this phosphorylation abrogated RocS binding to DNA. Overall, this study offers new insights into chromosome segregation in Streptococci and paves the way for a deeper understanding of RocS-like proteins in other bacteria.

**IMPORTANCE:** Bacteria have evolved a variety of mechanisms to properly segregate their genetic material during cell division. In this study, we performed a molecular dissection of the chromosome partitioning protein RocS, a pillar element of chromosome segregation in *S. pneumoniae* that is also generally conserved in the *Streptococcaceae* family. Our systematic investigation shed light on the molecular features required for successful pneumococcal chromosome segregation and the regulation of RocS by phosphorylation. In addition, our study also revealed that RocS shares functional domains with the Par protein, involved in an atypical plasmid segregation system. Therefore, we expect that our findings may serve to extend our understanding of RocS and RocS-like proteins, while broadening the repertoire of partitioning systems used in bacteria.

## INTRODUCTION

Chromosome inheritance is crucial in all domains of life. Unlike eukaryotes, bacteria do not rely on a spindle-like mitotic apparatus to organize chromosome segregation. Instead, they combine passive and active mechanisms to concomitantly replicate, segregate and transcribe their genetic material in synchronization with cell division (1). Most of our understanding of bacterial chromosome segregation comes from observations in the model bacteria *Bacillus subtilis*, *Escherichia coli* and *Caulobacter crescentus* (2). Although these studies have pointed towards shared mechanisms, they have also highlighted divergences, such as the relative importance of conserved mechanisms depending on the studied organism (3), as well as unique strategies to cope with various constraints specific to their physiology, mode of growth and environmental niches (4, 5).

One example of this duality is the broadly distributed ParABS system, responsible for the early segregation of the origin region through the combination of a CTPase (ParB) and an ATPase (ParA) activities (6). *C. crescentus*, in particular, relies on this system to orchestrate chromosome segregation (7). However, the system is absent in *E. coli* (8), while *B. subtilis* uses it mainly during sporulation (9). The versatility of the ParABS system is also illustrated by its plasmid-encoded version, which assists the segregation of low-copy plasmids to ensure their maintenance (8). Another ubiquitous factor for chromosome segregation is the Structural Maintenance of Chromosome (SMC) complex, SMC-ScpAB or its MukBEF and MksBEF counterparts (3). These complexes are proposed to participate in chromosome segregation via an ATP-driven loop-extrusion mechanism (2). Again, their importance varies, from having little impact on chromosome segregation in *C. crescentus* to being essential in *B. subtilis* and *E. coli* when grown in nutrient-rich medium (3).

Over the past decade, *Streptococcus pneumoniae* has emerged as a model organism for deciphering the molecular mechanisms that control the bacterial cell cycle (10). In particular, these studies offered crucial insights into the mechanisms of cell division and morphogenesis, pioneering the discovery that protein phosphorylation is a fundamental regulatory mechanism in the pneumococcal cell cycle (11–14). However, a fine understanding of chromosome segregation in this bacterium remains largely elusive. The first studies reported the presence of an incomplete ParABS system, in which the ParA protein was lacking (10, 15). Furthermore, this system, as well as the SMC complex, appeared only moderately involved in chromosome segregation, as their single and dual deletion only generated a low percentage of anucleate cells (15, 16). This suggested the existence of an additional player, which was identified recently as the Regulator of Chromosome Segregation (RocS) protein (17). Indeed, the deletion of *rocS* is extremely deleterious for chromosome segregation and is synthetic lethal with the deletion of *parB* and *smc*, thus establishing it as the main driver of chromosome segregation in *S. pneumoniae* (17). RocS is a protein bound to the membrane and the nucleoid *via* its C-terminal amphipathic helix (AH) and its N-terminal DNA-binding domain (DBD) respectively. Both domains are essential for RocS localization and function in chromosome segregation (17). However, the molecular mechanisms involved remain elusive. In this work, we performed a mutational dissection of RocS to reveal the key features required for chromosome segregation. In particular, we demonstrate that the coiled-coil domain (CCD) of RocS, which separates the AH and the DBD, can be divided into two subdomains, each serving distinct functions. . Furthermore, we revealed that a domain of unknown function (DUF 536) plays a major role in the RocS-assisted chromosome segregation. Finally, we show that the DBD is phosphorylated by StkP, thus negatively modulating DNA-binding and subsequent chromosome segregation. Overall, this study provides new insights into chromosome segregation in Streptococci and paves the way for a better understanding of RocS-related proteins in other bacteria.

## RESULTS

### The amphipathic helix of RocS requires multivalency to associate with the membrane

It was previously demonstrated that RocS harbors a C-terminal amphipathic helix (AH). This AH mediates the interaction of RocS with the membrane, which is crucial for its localization and function (17). To gain insights into the binding properties of the AH, we fused it to the C-terminus of mGFP and expressed the GFP-AH_RocS_ fusion ectopically under the control of an inducible promoter. We observed the fluorescence distribution within these cells using phase-contrast and fluorescence microscopy. The signal of GFP-AH_RocS_ was diffuse in the cytoplasm, indicating that a single AH was not sufficient to mediate a stable association with the membrane (Fig. 1A). We have reported previously that the AH of RocS shares a homologous sequence with the AH of MinD from *E. coli* (17) (Fig. S1). This protein is known to interact with the membrane via its AH upon ATP-dependent dimerization (18, 19). Therefore, we tested whether duplicating the AH of RocS would be sufficient to target the GFP to the membrane. In contrast to the monovalent construct GFP-AH_RocS_ and consistent with previous results obtained with the AH of MinD (19), the signal of GFP-(AH_RocS_)_2_ was mainly localized at the cell periphery (Fig. 1A). This indicates that this fusion binds to the membrane of *S. pneumoniae* and suggests that RocS requires at least dimerization to associate with the membrane.

**Figure 1:**
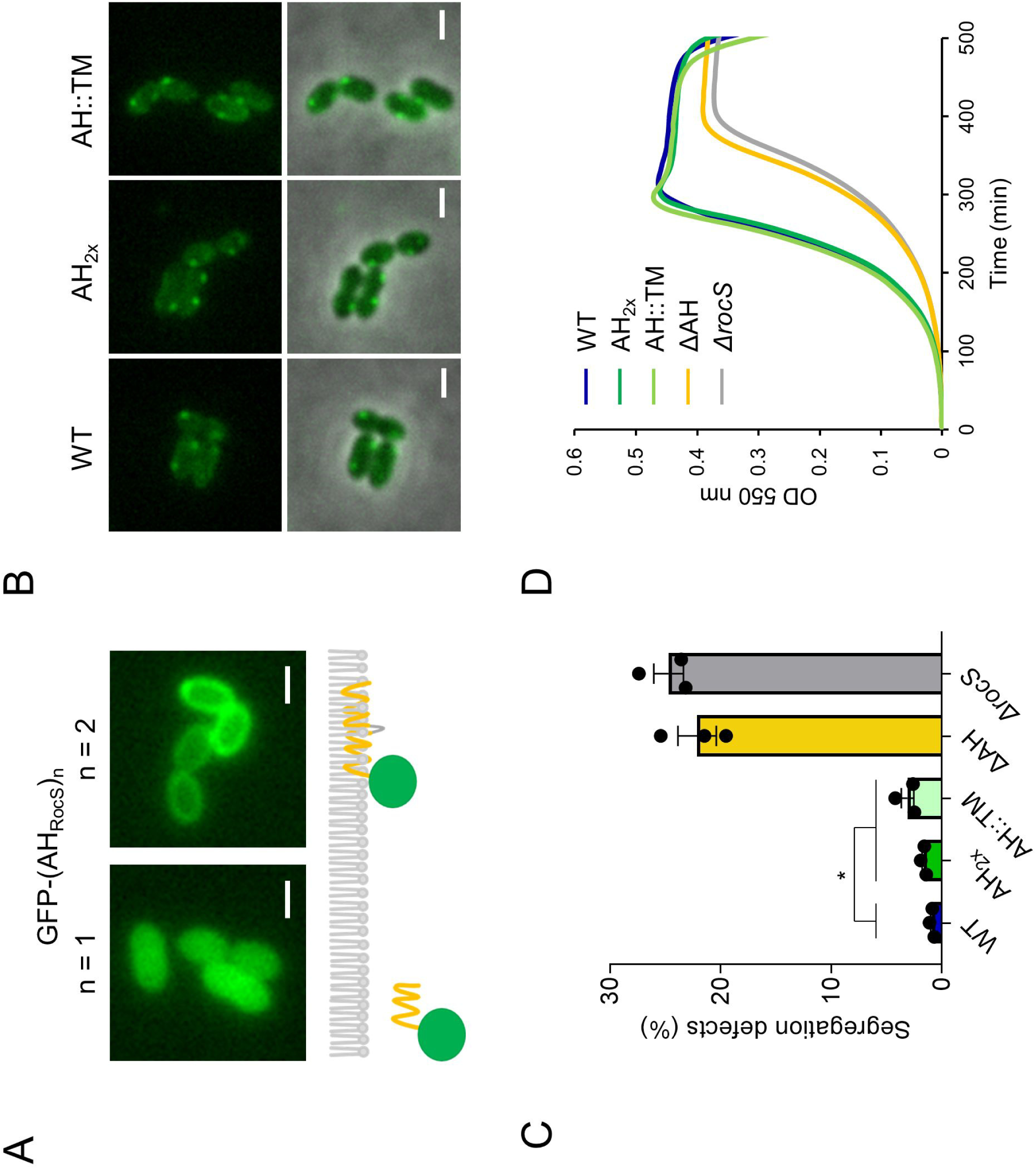
RocS has a MinD-like AH. (A) Spatial localization of a GFP probe fused to 1 or 2 AHs. Upper panel: representative fluorescence images of the *S. pneumoniae* R6 *bgaA::pComX-gfp-(AH_RocS_)_n_* strains with n = 1 (left) or 2 (right). Lower panel: schematic depiction of the localization of the probes. The single or the double AHs are represented in yellow, the GFP in green and the inner leaflet of the membrane in grey. (B) Representative images showing the localization of GFP-RocS WT and specified mutants. From top to bottom: GFP channel, overlay GFP and phase-contrast channels. Scale bar, 1 µm. (C) Bar chart representing the percentage of chromosome segregation defects observed in the *S. pneumoniae* R6 *rocS::rocS mutants ; (hlpA+) hlpA-mKate2* strain. Bars represent the mean (± SEM) of three independent experiments. The WT was compared separately with the RocS-AH_2x_ or the RocS-AH::TM mutants with an unpaired t-test (p-value 0.0167 and 0.0185 respectively). (D) Growth curves of *S. pneumoniae* R6 *rocS::rocS mutants* (mean of 3 biological replicates).

To investigate the dynamics of membrane association and its role in RocS function, we assessed the localization of GFP-tagged RocS harboring a duplicated AH (GFP-RocS-AH_2x_) expressed at the native chromosomal locus. Throughout the study, we confirmed the expression and stability of all our constructs with a polyclonal anti-GFP antibody (Fig. S2). The localization of GFP-RocS-_AH2x_ was similar to GFP-RocS WT, i.e. they formed bright foci close to the membrane (Fig. 1B). We also examined chromosome segregation using the merodiploid HlpA-mKate2 fluorescent reporter to visualize the nucleoid and quantify segregation defects (see Material and Methods) (16). Cells expressing RocS-AH_2x_ exhibited limited segregation defects (Fig. 1C, segregation defects RocS WT: 0.92%, N = 1,220 *vs* RocS-AH_2x_: 1.66%, N = 1,730 – p-value = 0.0167) and cell growth was not impacted (Fig. 1D). A similar phenotype was observed when we substituted the AH of RocS for the transmembrane domain of MapZ (TM-Ex_1_), a transmembrane protein unrelated to chromosome segregation (20–22). In particular, GFP-RocS-AH::TM displayed a WT-like localization (Fig. 1B) and its untagged version supported normal growth (Fig. 1D). Chromosome segregation was also slightly impacted compared to the WT (Fig. 1C, segregation defects RocS WT: 0.92%, N = 1,220 *vs* RocS-AH::TM: 3.13%, N = 1,168 – p-vlaue = 0.0185). Finally, we compared both mutants to a strain expressing a version of RocS devoid of AH (RocS-ΔAH) which has been previously characterized (17). As expected, this mutant was considerably more impacted, with growth and chromosome segregation defects comparable to the markerless deletion mutant *ΔrocS* (Fig. 1C and 1D, segregation defects RocS-ΔAH: 22.16%, N = 1,193 *vs ΔrocS*: 24.73%, N = 1,077 – p-value = 0.3067). In summary, RocS requires prior oligomerization to interact with the membrane and this interaction is required for RocS function. Additionally, altering the nature of the membrane anchor does not drastically impair RocS function.

### The coiled-coil domain drives RocS oligomerization

Since RocS needs to oligomerize to interact with the membrane, we investigated its ability to self-interact using a bacterial adenylate cyclase-based two-hybrid (BACTH) assay. Specifically, the N-terminus of RocS and derivatives was fused to either the T18 or the T25 fragment of the catalytic domain of adenylate cyclase (CyaA) from *Bordetella pertussis* (23).

As anticipated, we detected a strong interaction between full-length CyaA-RocS hybrids after 48h of incubation (Fig. 2A).

**Figure 2:**
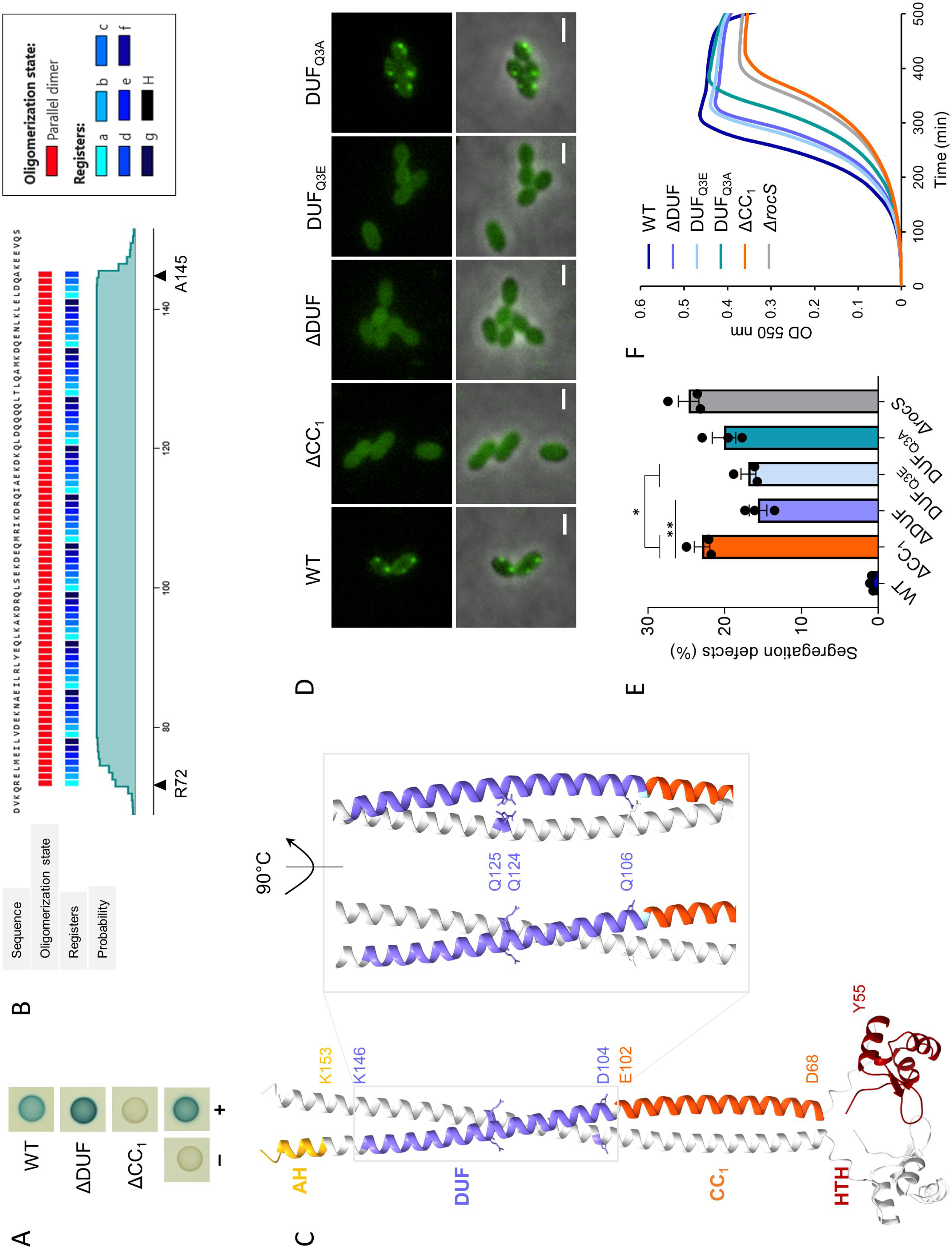
The conserved DUF is a major determinant of RocS function. (A) The self-interaction between RocS, RocS-ΔDUF and RocS-ΔCC_1_ were assessed by bacterial two hybrid assay. A blue coloration indicates a positive interaction as depicted by the positive control (+) and the negative control (-) (B) Coiled-coil prediction by the bioinformatic tool CoCoNat. The sequence of full-length RocS was submitted for analysis. The arrows indicate the bounded residues of the unique predicted CCD. (C) AlphaFold3 model of a RocS dimer. Chain A is depicted as a grey ribbon while chain B is colored according to the corresponding domain (dark red 1-55: HTH, orange 68-102: CC_1_, violet 104-146: DUF, yellow 153-163: AH. The bounded residues of each domain and the 3 conserved glutamines of the DUF are annotated. The 3 conserved glutamines are represented as amino acids to visualize their side chain. A magnified view of the DUF at 0°C and 90°C angles enables a better visualization of the conserved glutamines. (D) Representative images showing the localization of GFP-RocS WT and specified mutants. The GFP channel and the overlay between the GFP and the phase-contrast channels are shown. Scale bar, 1 µm. (E) Bar chart representing the percentage of chromosome segregation defects observed in the *S. pneumoniae* R6 *rocS::rocS mutants ; (hlpA+) hlpA-mKate2* strains. Bars represent the mean (± SEM) of three independent experiments. Statistical comparisons between the RocS-ΔCC1 mutant and the RocS-ΔDUF or the RocS-DUFQ3E mutant were performed with an unpaired t-test (p-value 0.0089 and 0.0128 respectively) (F) Growth curves of the WT and specified mutants (mean of 3 biological replicates).

We next sought to identify the determinants of this self-interaction. According to two *in silico* coiled-coil structure prediction tools (CoCoNat (24) and CoCoPRED (25)), the C-terminal AH of RocS is immediately followed by a coiled-coil domain (CCD) spanning approximately the entire length of the helix joining the AH to the DBD (Fig. 2B and S3). The CCD is predicted to assemble into a parallel dimer (Fig. 2B and S3). Additionally, the analysis of the sequence indicates that the CCD contains a conserved domain of unknown function DUF 536 (Pfam PF04394, InterPro IPR007489), recently renamed RocS-like_C and herein referred to as DUF. The CCD was thus divided into two subdomains, the CC_1_ from residues D68-E102 and the DUF from residues D104-D146 (Fig. 2C), which were examined separately to decipher their potential contribution to the oligomerization of RocS. Removing CC_1_ completely abrogated the interaction between the CyaA fusions (Fig. 2A). In line with this observation, GFP-RocS-ΔCC_1_ displayed a signal dispersed in the cytoplasm and no longer formed distinct foci, nor was enriched at the membrane (Fig. 2D). Additionally, a strain expressing RocS-ΔCC_1_ was characterized by severe growth and chromosome segregation defects comparable to a *ΔrocS* mutant (Fig. 2E and 2F), thus confirming that self-interaction is crucial for the function of RocS. By contrast, we detected a strong interaction between CyaA-RocS-ΔDUF derivatives (Fig. 2A), indicating that this domain is not required for homotypic interaction. However, GFP-RocS-ΔDUF exhibited a signal similar to the ΔCC_1_ fusion, i.e. cytosolic (Fig. 2D), revealing that this domain is still essential for the interaction of RocS with the membrane. Surprisingly, the expression of the corresponding untagged truncated protein resulted in intermediate growth and chromosome segregation defects (Fig. 2E et F). Indeed, while the deletion of RocS or the ablation of the CC_1_ domain generated around 24% of cells with an abnormal chromosome content (Fig. 2E, segregation defects *ΔrocS*: 24.73%, N = 1,077 *vs* RocS-ΔCC_1_: 22.97%, N = 1,318 – p-value = 0.3564), the absence of the DUF only resulted in 16% of cells with a chromosome segregation defect (Fig. 2E, segregation defects RocS-ΔDUF: 15.69%, N = 1,915 *vs ΔrocS*: p-value = 0.0068 ; *vs* RocS-ΔCC_1_: p-value = 0.0089). Collectively, these results highlight two distinct modules within the CCD, the CC1 and the DUF, each contributing differently to RocS function. While the CC_1_ is required for the homodimerization and subsequent membrane association of RocS, the DUF likely serves another function that is crucial for RocS-mediated chromosome segregation.

### Three conserved glutamines underpin the function of the DUF 536

To build upon our observations regarding the DUF, we analyzed its sequence conservation. According to Pfam (26), the DUF536 domain is defined by the presence of three highly conserved glutamines (Fig. 2C, S4). To test whether these residues contribute to the function of RocS, we mutated them into alanine (DUF_Q3A_) or glutamate (DUF_Q3E_) and investigated the localization of their cognate GFP fusion. First, we observed that the signal displayed by GFP-RocS_Q3E_ was cytosolic, reminiscent of the localization of GFP-RocS-ΔDUF (Fig. 2D). Furthermore, the RocS-DUF_Q3E_ mutant phenocopied the RocS-ΔDUF mutant, presenting similar growth and segregation defects (Fig. 2E and F, segregation defects RocS-DUF_Q3E_: 16.93%, N = 1,172 *vs* RocS-ΔDUF: 15.69%, N = 1,915 – p-value = 0.4532). Unexpectedly, we observed a completely different behavior when the glutamines were mutated into alanines. Indeed, GFP-RocS-DUF_Q3A_ formed bright static foci, characteristic of the localization of the WT fusion (Fig. 2D). However, the expression of RocS-DUF_Q3A_ did not result in a WT-like phenotype. On the contrary, growth was even more affected than for a RocS-DUF_Q3E_ mutant and the segregation defects were also very severe (Fig. 2E and F, segregation defects RocS-DUF_Q3A_: 20.10%, N = 1,002 *vs* RocS-DUF_Q3E_: p-value = 0.1551). In summary, these results establish the conserved glutamines as major determinants of the function of the DUF. Moreover, they reveal the previously overlooked importance of this domain for RocS function.

### The DNA-binding domain of RocS is phosphorylated by StkP

We have previously shown that the first 55 amino acids of RocS are predicted to form a helix-turn-helix (HTH) domain, a DNA-recognition motif in which two α-helices are separated by a sharp turn (17, 27). An AlphaFold3 prediction further suggests that the HTH of RocS could be followed by an additional β-strand hairpin (wing), characteristic of the winged HTH (wHTH) subfamily (Fig. 2C) (27). Interestingly, a DALI search of the wHTH prediction model of RocS revealed that this domain shares structural homology with the *Staphylococcus aureus* plasmid partitioning protein Par (28) and the *Bacillus subtilis* chromosome segregation protein RacA (4, 29) (Fig. 3A), which both harbor a MerR-type wHTH domain (30, 31). According to the structure of the corresponding protein-DNA complexes, the helix α2 is inserted in the major groove while the wing primarily contacts the phosphate backbone and possibly the minor groove (30, 31). This interaction is also suggested by the Alphafold3 modeling of a RocS dimer in complex with a double-stranded (ds) DNA fragment (Fig. 3B).

**Figure 3:**
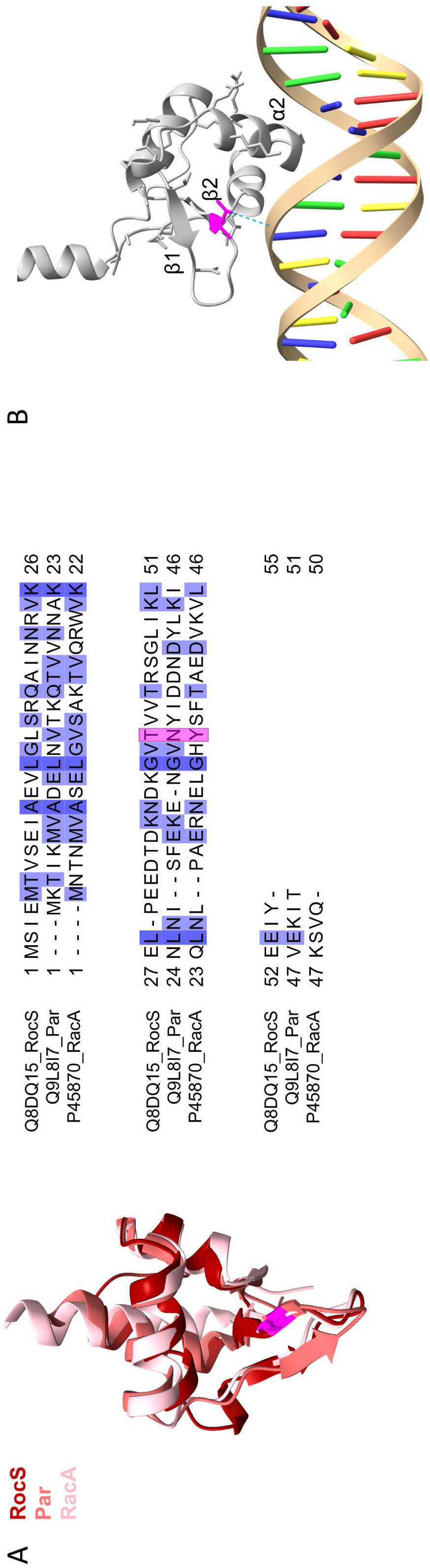
RocS is phosphorylated on threonine 41. (A) Structure and sequence alignment of the wHTH domains of RocS, Par and RacA. Left panel: superimposition of the AlphaFold3 model of the wHTH of RocS with the structures of Par (8csh-A) and RacA (5i44-B) after a pairwise alignment using the MatchMaker tool in ChimeraX with default settings. Across all 50 pairs RocS-Par, RMSD between 35 pruned atom pairs was 0.942 angströms. Across all 61 pairs RocS-RacA, RMSD between 35 pruned atom pairs was 1.024 angströms. T41 (from RocS), Y36 (from RacA), N36 (from Par) are highlighted in pink. Right panel: sequence alignment of the wHTH of RocS with the wHTH of Par and RacA taking into account their secondary structures (53). Sequences are indicated with UniProt identifiers, followed by the protein name. The alignment is colored according to the standard percentage identity color scheme as implemented in Jalview (https://www.jalview.org/). T41 from RocS and the corresponding residues from Par (N36) and RacA (Y36) are highlighted in pink. (B) AlphaFold3-generated model of a RocS dimer in complex with DNA (zoom in on one chain, depicted as a grey ribbon). The DNA-recognition helix and the wing are annotated as α2 and β1-β2 respectively. T41 is highlighted in pink and a putative hydrogen bond between the hydroxyl side-chain and the phosphate backbone of DNA is depicted as a blue dashed line.

Interestingly, the threonine 41 (T41), located within the wing (Fig. 3A and 3B), was recently found to be phosphorylated when cells were grown in the presence of ampicillin (14). This antibiotic is known to increase the activity of the unique Serine-Threonine kinase StkP in *S. pneumoniae* (12). To assess whether T41 is also phosphorylated by StkP in normal growth conditions, we immunoprecipitated a GFP-RocS fusion expressed from the native promoter at the chromosomal locus in a WT or a *ΔstkP* genetic background and performed a phosphoproteomic analysis following a phosphopeptide enrichment procedure (see Materials and Methods). We detected that T41 was phosphorylated in WT cells but not in the strain *ΔstkP* (Fig. S5), thus confirming that RocS is a *bona fide* substrate of StkP in normal growth conditions.

### Modulation of DNA-binding by phosphorylation affects chromosome segregation

The structural alignment of the wHTH domains from RocS and RacA shows that T41 corresponds to Y36 in RacA (Fig. 3A), an amino acid previously implicated in the interaction of RacA with the phosphate backbone of DNA (30). Therefore, we hypothesized that phosphorylation might affect its interaction with DNA. To explore this idea, we overproduced and purified a phosphomimetic (T41D) version of RocS from *E. coli* (Fig. S5). As previously described (17), this construct did not contain the AH and was labeled with a 6xHistidine tag, which was subsequently removed. The binding of the purified proteins to DNA was then assayed by an electrophoretic mobility shift assay (EMSA). RocS WT-ΔAH and RocS G15P-ΔAH (Fig. 4A), previously shown to respectively retain or lose their DNA-binding ability (17), were respectively used as positive and negative controls. We observed that DNA incubated with RocS T41D-ΔAH migrated freely, suggesting that this mutant was unable to interact stably with DNA (Fig. 3C). This observation was confirmed by measuring the binding affinities of RocS-WT and RocS-T41D using microscale thermophoresis (MST). Indeed, the affinity of RocS-T41D for DNA was considerably decreased with an affinity constant (K_D_) above 500 µM, more than ten times higher than the WT (K_D_ = 34 µM) (Figure 4B). This indicates that T41 is directly involved in the interaction with DNA and further suggests that the phosphorylation of T41 could disrupt this interaction.

**Figure 4:**
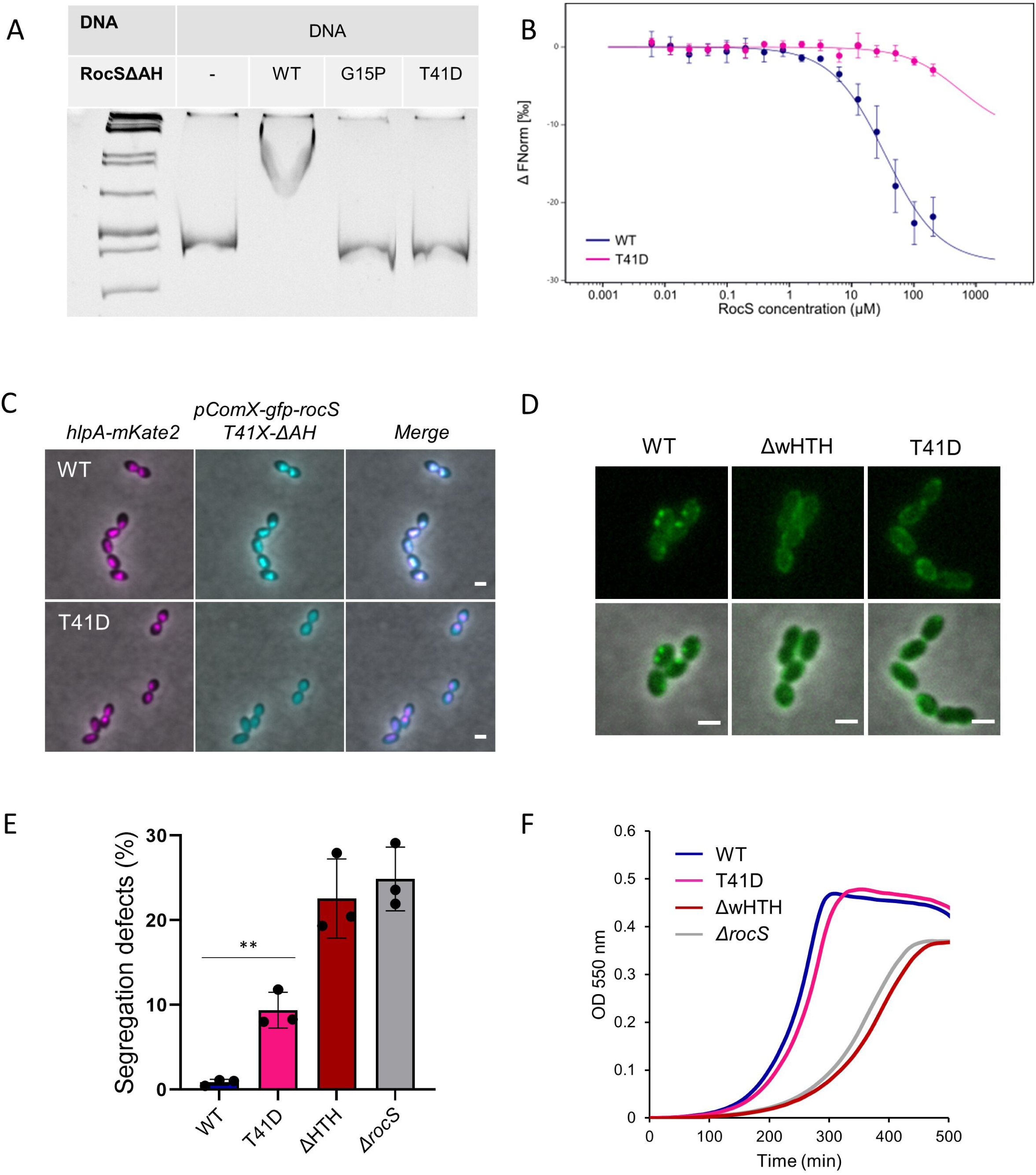
RocS phosphorylation and DNA-binding and functionality *in vivo*. (A) Electrophoretic mobility shift assay of RocS-ΔAH (WT and point mutants) with a 500 bp ds DNA fragment. From the left to right lane: GeneRuler 1kb DNA ladder (Thermo Scientific^TM^), free DNA, DNA and RocS WT-ΔAH, DNA and RocS G15P-ΔAH, DNA and RocS T41D-ΔAH. (B) Microscale thermophoresis binding assay of dnaC_Cy5 to increasing concentrations of RocS WT-ΔAH (blue) or RocS T41D-ΔAH (pink). The variation of the normalized fluorescence (ΔFNorm) is plotted against RocS concentration. Measurements are represented by dots (mean of three independent experiments) and the fitted curve by a line. The error bars represent the standard deviation. (C) Co-localization of GFP-RocS WT or T41D-ΔAH. Representative fluorescence and phase-contrast images of the *S. pneumoniae* R6 *bgaA::pComX-gfp-RocS WT-ΔAH, (hlpA+) hlpA-mKate2* and R6 *bgaA::pComX-gfp-RocS T41D-ΔAH, (hlpA+) hlpA-mKate2* strains. From left to right, the mCherry and phase-contrast channels overlay, the GFP and phase-contrast channels overlay and the mCherry, the GFP and phase-contrast channels overlay are shown. Scale bar, 1 µm. (D) Representative images of the localization of GFP-RocS WT and specified mutants. The GFP channel and the overlay between the GFP and the phase-contrast channels are shown. Scale bar, 1 µm. (E) Bar chart representing the percentage of chromosome segregation defects observed in the *S. pneumoniae* R6 *rocS::rocS mutants ; (hlpA+) hlpA-mKate2* strain. Bars represent the mean (± SEM) of three independent experiments. Statistical comparison between the WT and RocS T41D strain was performed using an unpaired t-test (p-value = 0.0024). (F) Growth curves of the WT and the specified mutants (mean of three biological replicates)

We also examined the nucleoid-binding properties of RocS T41D *in cellulo*. For this purpose, we constructed a fusion of GFP with a truncated version of RocS, in which the AH domain was removed, and expressed it ectopically under the control of an inducible promoter. As observed previously when expressed from the native chromosomal locus (17), GFP-RocS-ΔAH co-localized with the nucleoid when expressed from an inducible promoter (Fig. 4C). By contrast, its phosphomimetic counterpart GFP-RocS-T41D-ΔAH no longer co-localized with the nucleoid and displayed a uniform signal inside the cytoplasm (Fig. 4C). Taken together, these results show that mimicking phosphorylation impairs DNA binding. This further suggests that phosphorylation negatively modulates the ability of RocS to bind DNA. Finally, we interrogated the functionality of the phosphomimetic mutant *in vivo* by looking at its subcellular localization and its effect on chromosome segregation and growth. As shown in Figure 4D, the GFP-RocS T41D fusion did not form the bright foci characteristic of the wild-type and displayed a membrane-enriched localization instead, similar to a truncated mutant lacking the wHTH (GFP-RocS-ΔwHTH, Fig. 4D). As self-interaction and subsequent membrane association is not affected in both cases (Fig. S7 and 4D), this signifies that RocS does not form higher-order assemblies in the absence of the DBD or when it is phosphorylated. Yet, RocS T41D and RocS-ΔwHTH mutants exhibited a strikingly different phenotype. While the expression of RocS-ΔwHTH resulted in a level of chromosome segregation defects comparable to a *ΔrocS* mutant (Fig. 4E, segregation defects RocS-ΔwHTH: 22.57%, N = 1,395 *vs ΔrocS*: 24.88%, N = 1,511 – p-value = 0.5416), the expression of RocS T41D generated two times less chromosome segregation defects (Fig. 3F, segregation defects RocS T41D: 9.37%, N = 1,733). Moreover, the phosphomimetic mutant presented a normal growth, which contrasted with the strong growth defect observed in both RocS-ΔwHTH and *ΔrocS* mutants (Fig. 4F). In conclusion, the phospho-regulation of RocS is less deleterious than the removal of its DBD.

## DISCUSSION

Chromosome segregation in bacteria is a critical process that ensures the faithful partitioning of the genetic material to each daughter cell. Although *S. pneumoniae* encodes for the well characterized proteins ParB and SMC, which are critical for chromosome segregation in other bacteria (3, 6), these two proteins appear to have a minor contribution (15, 16). In addition, the pneumococcus lacks ParA, the P-loop ATPase that directs segregation to opposite sides of the cell (10, 15). In this context, the discovery of RocS as the major contributor to chromosome segregation in *S. pneumoniae* revealed that the molecular mechanisms governing chromosome segregation are significantly different from those at play in other bacteria (17). In this study, we have dissected the molecular features of RocS to shed light on this alternative partition mechanism and understand the relative contribution of each domains of RocS to chromosome segregation.

Our results show that RocS is organized into three major functional domains: a C-terminal AH acting as a membrane anchor, a central CCD mediating multimerization and an N-terminal wHTH domain binding DNA. Although previously classified as a MarR-type wHTH domain (17), advances in secondary structure prediction (32) reveal that the DBD is more likely to have a MerR-type wHTH fold (Fig. 3A and B). In particular, the RocS wHTH shows strong structural similarity to two distant members of the MerR-type wHTH family: the *B. subtilis* chromosome segregation protein RacA (29, 30) and the *S. aureus* plasmid partitioning protein Par (28, 31) (Fig. 3A). Both proteins contain three domains: an N-terminal wHTH domain, a central coiled-coil region, involved in oligomerization and possibly protein-protein interactions, and a C-terminal disordered domain (30, 31). RacA contributes to DNA segregation during sporulation through the binding of repetitive centromere-like sequences on the nucleoid and the interaction with the polar anchor DivIVA (4, 29, 30, 33). Therefore, the RacA protein can be considered as a sub-cellular positioning system that tethers the nucleoid to the pole. However, the biased localization of RocS at the cell equators (17) argue against the implication of DivIVA or an alternative polar anchor in RocS-dependent DNA segregation. On the other hand, Par is a single protein partition system involved in the segregation of the low-copy number plasmid pSK1 (28). Interestingly, the architecture of Par shares an additional similarity with RocS, as it is also predicted to contain the DUF 536. Moreover, an alignment of both sequences reveals that the C-terminal amino acid-stretch of Par is practically identical to the AH of RocS and MinD (Fig. S1).

The conservation of the DUF-AH module in Par (Fig. S1) suggests that it could also bind to the membrane and share a common DNA partition mechanism with RocS. While more work is required to identify such mechanism, it can narrow-down previously suggested scenarios (31). By analogy with the partitioning systems of eukaryotic extrachromosomal elements (34), it has been proposed that the Par-dependent plasmid segregation system could bind both specific sequence repeats on the plasmid and degenerated sequence repeats on the chromosome, thus using the nucleoid as a vehicle (31). Therefore, the Par protein could be considered as a sub-cellular positioning system that tethers plasmids to the nucleoid. However, this hitchhiking model cannot be applied to RocS and direct evidence for this mechanism in Par is still lacking. Alternatively, we propose that RocS, and possibly Par, could be considered as a sub-cellular positioning system that tethers the targeted DNA to the membrane. However, this still poses the question of how a single DNA-binding protein can drive segregation by membrane tethering.

The absence of an energy-driven partitioning system might indicate a predominant role for entropy-driven segregation. Entropic forces have previously been proposed to drive bacterial chromosome segregation by favoring the natural separation of replicated DNA molecules, thereby increasing the overall disorder within the cell (35, 36). However, purely entropic forces are generally insufficient for bacterial chromosomes segregation and could even hinder chromosome segregation if precise physical conditions are not met (37). In this context, the proteins previously identified as segregation factors could help create conditions that promote entropy-driven segregation, ensuring that each daughter cell receives one copy of the chromosome. For instance, a biophysical model suggests that ParB and SMC alter the effective topology of the chromosome, thereby redirecting and amplifying entropic forces to enable accurate chromosome segregation (37). Consistently, by tethering the nucleoid to the membrane, RocS could alter the physical state of the chromosome to create the right conditions for entropy-directed segregation. As previously mentioned, the single or double deletion of *parB* and/or *smc* only generates a small percentage of anucleate cells in *S. pneumoniae*. However, the deletion of either gene is synthetic lethal when combined with the deletion of *rocS* (17). Altogether this suggests that ParB, SMC and RocS may have additive and possibly synergetic actions on chromosome segregation by altering the topology of the chromosome.

An alternative way to drive the chromosome segregation would be through the intervention of another protein. As mentioned before, RocS and Par are predicted to contain the same domain of unknown function (DUF 536). This DUF domain is not required for RocS homodimerization (Fig. 2A) but is crucial for its function in chromosome segregation (Fig. 2E). Moreover, when we mutated three conserved glutamines of the DUF domain into alanines, we observed a strong effect on chromosome segregation while the localization of RocS remained unaffected (Fig. D and E). Since we demonstrated that the formation of bright foci close to the membrane, characteristic of the WT localization, depends on the ability of RocS to dimerize, associate with the membrane (Fig. 2D), and interact with DNA (Fig. 4D), the DUF domain, and particularly the three conserved glutamines, must serve another important function for RocS-mediated chromosome segregation. While we cannot rule out that these conserved glutamines may stabilize interactions at the dimer or oligomer interface, they might also be involved in interactions with an additional partner. Interestingly, a sequence alignment of all proteins predicted to contain the DUF indicates that most proteins also harbor a putative C-terminal MinD-like AH (Fig. S8). Therefore, it is tempting to speculate that the DUF 536 and the AH form a functional unit, which could work in tandem with a yet to be identified partner to form a complete partitioning system. Future work is definitely needed to address this question.

Finally, we showed that the N-terminal wHTH domain, which mediates DNA binding, is phosphorylated by StkP on T41 (Fig. S5). Specifically, we revealed that a phosphomimetic mutation (T41D) affects the interaction of RocS with DNA (Fig. 4A, B and C). Protein phosphorylation has been previously suggested to regulate chromosome segregation in mycobacteria (38) and possibly *B. subtilis* (39). On the other hand, Serine-Threonine phosphorylation by StkP in *S. pneumoniae* is generally considered as a central regulator of the pneumococcal cell elongation and division, in particular by modulating cell elongation, constriction and final separation (20, 40–42). Therefore, our study identifies an additional layer of regulation controlled by StkP, related to chromosome segregation. In sum, StkP would not only be dedicated to the control of cell division but would act as a comprehensive coordinator of several, if not all, processes required to achieve a successful cell cycle. This hypothesis is supported by the characterization of other potential substrates of StkP and/or its cognate phosphatase PhpP that are involved in DNA replication and repair such as MutL, SsB, DnaD and PolA (14, 43). Future studies focusing on the dynamics of RocS phosphorylation during the cell cycle, also in relation to the phosphorylation of the other substrates, should clarify the role of RocS phosphorylation and subsequent perturbation of its association with DNA during the cell cycle.

## MATERIALS AND METHODS

### Strains and growth conditions

Strains of *S. pneumoniae* were grown in C+Y or THY medium at 37°C. For growth on plate, THY agar supplemented with 3% horse blood was used. The expression of fluorescent probes under the control of the pComX promoter was achieved by addition of 1 μM of the inducer ComS (44) 30 minutes prior to imaging or for the whole culture duration for crude extracts preparation. Growth was monitored with a Tecan SUNRISE microtiter plate reader set up with a 550 nm filter. Cells were first grown up to OD_550_ = 0.3 in C+Y medium. They were then diluted at OD_550_ = 0.0001 and transferred in a Nunc Edge 96-well, non-treated, flat-bottom MicroplateGreen plate, whose lid was preventively treated with a 0.05% Triton-X100 and 20% ethanol mixture to avoid condensation. Growth was monitored at 37°C with OD measurements taken every 10 minutes after brief shaking. *E. coli* XL1-Blue, *E. coli* BL21 star (DE3) and *E. coli* BTH101 strains were used for cloning, recombinant protein expression and BACTH respectively. All were grown in LB medium supplemented with appropriate chemicals and antibiotics.

### Allelic replacement mutagenesis

The construction the strains used in this study is detailed in table S1. The oligonucleotides used for cloning purposes are listed in table S2. Strains of *S. pneumoniae* were derived from R800 rpsL1 (resistant to streptomycin) and transformed with PCR products of interest. Briefly, bacteria were cultivated in C+Y medium (pH 6.8) at 37°C without agitation and competence was induced at around OD_550_ = 0.1 by addition of the synthetic competence stimulating peptide 1 (CSP1) (45) at 100 ng/10^8^ cells. DNA was then mixed with competent cells and incubated at 37°C for 30 minutes before plating dilutions of the transformed bacteria mixed in THY-agar supplemented with 3% of defibrinated horse blood. After 2 h of incubation at 37°C, an additional THY-agar layer supplemented with the appropriate antibiotic (streptomycin 200 μg/mL, kanamycin 250 μg/mL or chloramphenicol 4.5 μg/mL) was poured and the cells were left to incubate overnight. Gene replacement or deletion was achieved by homologous recombination following a 2-step selection process (46). First, the gene was replaced by a *kanR-rpsL* cassette restoring sensitivity to streptomycin while conferring resistance to kanamycin. Colonies were selected on plates supplemented with kanamycin (0.25 mg/mL). The second step consisted in the cassette’s removal and selection on THY-agar plates supplemented with streptomycin (0.2 mg/mL). All strains were verified by colony PCR, antibiotic sensitivity testing and locus sequencing.

The merodiploid copy of *hlpA* fused to *mKate2* was inserted along with a chloramphenicol resistance marker as previously described (16) and colonies were selected on THY-agar plates supplemented with chloramphenicol (4.5 μg/mL).

### Plasmids construction

The construction of the plasmids used in this study is detailed in table S1. The oligonucleotides used for cloning purposes are listed in table S2. BACTH plasmids carrying the full-length RocS were constructed as follow: *rocS* was PCR-amplified from the *S. pneumoniae* R6 strain and inserted in the pUT18C and pkT25 vectors digested XbaI and Acc65I restriction enzymes (New England Biolabs). The derivatives deletion mutants were PCR amplified using designated primers and the resulting linear plasmids were re-circularized using Gibson assembly (New England Biolabs). Expression plasmids carrying mutated versions of RocS were PCR-amplified from a pt7-7 vector carrying *rocS*-*ΔAH-6His* (17) and circularized using Gibson assembly (New England Biolabs). Plasmids were verified by sequencing and stored in the *E. coli* XL1B strain.

### Microscopy techniques and image analyses

The cells were grown up to OD_550_ = 0.1-0.2, concentrated 10 times by centrifugation and resuspension, and 1 μL was spotted onto a 1% agarose / C+Y pad. Visualization was performed using a Nikon TiE microscope fitted with an Orca-CMOS Flash4 V2 camera with a 100 x 1.45 objective. Images were collected using NIS-Elements (Nikon) and were analyzed with ImageJ (http://rsb.info.nih.gov/ij/) and the MicrobeJ plugin (47). For the quantification of the chromosome segregation defects, cells were manually classified in the MicrobeJ editor according to their HlpA-mKate2 profile. Specifically, cells were considered as missegregated when exhibiting 1) no HlpA-mKate2 signal, 2) a cytosolic HlpA-mKate signal 3) an assymetrical HlpA-mKate2 distribution between two dividing cells. Statistical analyses were performed on biological triplicates, using unpaired Student’s *t*-test with MicrobeJ and GraphPad Prism 10 software (www.graphpad.com).

### Preparation of S. pneumoniae crude extracts and immunoblot analysis

The strains were grown in C+Y medium up to OD_550_ = 0.3-0.5, pelleted and resuspended in TE buffer (10 mM Tris/HCl, pH 8.0, 1mM EDTA) supplemented with a cocktail of protease inhibitors (CLAPA [Chymostatine 1 μg/mL, Leupeptine 1 μg/mL, Antipaine 1 μg/mL, Pepstatine 1 μg/mL, Aprotinine 5 μg/mL, Sigma-Aldrich] and 1 µg/mL cOmplete, EDTA-free, Roche). Cells were sonicated (Branson 450) with 20 pulses at 3/30 followed by 2 times 20 pulses at 4/40. The protein concentration was determined by the Bradford method using the Pierce protein assay reagent. 20 μg of total protein extract were loaded onto an SDS-PAGE gel and analyzed by western blotting. A polyclonal anti-gfp primary antibody (Amsbio, TP401) diluted in 1% BSA-TBST (1:10,000) was used in combination with a goat anti-rabbit secondary antibody coupled with horseradish peroxidase (BioRad, 170-6515) diluted in 1% BSA-TBST (1:5,000). The signal was revealed using a SuperSignal^TM^ West Pico PLUS Chemiluminescent Substrate kit (Thermo Scientific^TM^, 34580) according to the manufacturer’s instructions.

### Bacterial adenylate cyclase two hybrid (BACTH) assay

The BACTH assay was perfomed with the BACTH system kit (Euromedex) according to the manufacturer’s protocol (48). Briefly, N-terminal fusions of RocS full-length or truncation mutants with T18 or T25 were co-transformed and re-streaked on an LB-agar plate supplemented with ampicillin 100 μg/mL, kanamycin 50 μg/mL, isopropyl-β-D-thiogalactopyranoside (IPTG) 0.5 mM, and X-gal 100 µg/mL. Plates were grown at room temperature and photos were taken at 24, 48 and 72 hours to monitor the color shift of the colonies.

### Protein production and purification

Recombinant plasmids overexpressing RocS-ΔAH-6His and its variants were transformed *E. coli* BL21 STAR. Cells were grown in 2 L of LB medium supplemented with ampicillin (100 µg/mL) up to OD_600_ = 0.4-0.6 before inducing protein expression with 0.5 mM of isopropyl β-D-1-thiogalactopyranoside (IPTG) and further incubation at 25°C with shaking overnight.

Cells were collected by centrifugation and resuspended in buffer A (25 mM Tris/HCl, pH 7.5, 1 M NaCl, 10% glycerol) supplemented with a cocktail of protease inhibitors (CLAPA), 6 µg/mL of DNase I (Roche), 6 µg/mL of RNase A (Roche) and 10 µg/mL of lysozyme (Sigma-Aldrich). After 3 passages through a Continuous Flow Cell Disruptor (Constant Systems Ltd) at 1.5 kbar, membranes were further solubilized by the addition of 1% Triton (Euromedex) for 30 minutes. The lysate was clarified via centrifugation (14, 000 x g, 30 min, 4°C) and the supernatant was diluted with buffer A1 (25 mM Tris/HCl, pH 7.5 ; 10 % glycerol) to lower the concentration of NaCl to 500 mM (or 300 mM for RocS G15P-ΔAH-6His) before loading onto a 5 mL HisTrap column (Cytiva). The column was washed with 10% of buffer B (25 mM Tris/HCl, 500 mM NaCl, 300 mM imidazole, 10% glycerol) before being eluted following a linear gradient. Peak fractions were pooled and dialyzed overnight at 4°C in the presence of Tobacco Etch Virus (TEV) protease (0.025 mg per mg of protein), 1 mM DTT and 0.5 mM EDTA against buffer A2 (25 mM Tris/HCl, 300 mM NaCl, 10% glycerol) supplemented with 1 mM DTT and 0.5 mM EDTA. The next day, the cleaved sample was loaded onto a 5mL HisTrap column (Cytiva). The flow-through was collected, concentrated and injected onto a HiLoad 16/600 Superdex 200 pg size exclusion chromatography column (Cytiva) equilibrated in buffer C (25 mM Tris/HCl, pH 7.5, 150 mM NaCl, 5% glycerol). Peak fractions were pooled, concentrated up to 10-20 mg/mL and flash frozen in liquid nitrogen for subsequent storage at -70°C.

### Immunoprecipitation and mass spectrometry analysis

*S. pneumoniae* strains carrying a chromosomally encoded GFP-tagged RocS in a WT or Δ*stkP* backgrounds were grown at 37°C in 500 mL of THY broth until OD_550_ = 0.4. Cells were pelleted and washed with buffer A (20 mM Tris/HCl pH 7.5, 200 mM NaCl). The pellet was resuspended in buffer B (100 mM Tris/HCl, 150 mM NaCl, 5 mM MgCl_2_) supplemented with a cocktail of protease inhibitors (CLAPA and 1 µg/mL cOmplete, EDTA-free, Roche) and phosphatase inhibitors (Phosphatase Inhibitor Cocktail 2, Sigma-Aldrich), 6 µg/mL of DNase I (Roche), 6 µg/mL of RNase A (Roche), 50 U of mutanolysine (Sigma-Aldrich) and 8 mg/mL of lysozyme (Sigma-Aldrich). The cells were then sonicated (Branson 450) at 4°C for 4 x 1 minute at 4/40. Lysates were further incubated with 1% Triton-X100 (Euromedex) for 1h at 4°C, then cleared by centrifugation for 30 minutes at 14,000 x g. The lysates were incubated with 40 µL of GFP-TRAP® agarose resin (Chromotek) for 2 hours at 4°C on a rotating wheel. After collection by centrifugation, the beads were washed 3 times with buffer C (10 mM Tris/Hcl, pH 7.5, 150 mM NaCl, 500 µM EDTA) containing protease and phosphatase inhibitors. Finally, GFP-RocS was eluted with 30µL of Laemmli 2X and heating at 95°C for 10 minutes. The eluted samples were run onto a SDS-PAGE gel 12.5 % and stained with Coomassie.

### Nano LC-MS/MS analysis of purified GFP-RocS

Protein sample in gel was excised from SDS-PAGE and subjected to manual in-gel reduction, alkylation and digestion. The gel band was reduced with 10 mM DTT in 100 mM NH_4_HCO_3_ (Sigma Aldrich) for 1 h at 57°C and alkylated for 1h in the dark with 55 mM iodoacetamide in 100 mM NH_4_HCO_3_ (Sigma Aldrich), washed in 25 mM NH_4_HCO_3_, dehydrated with acetonitrile and dried in a speed-vac. Then the gel pieces were rehydrated with 40 µL trypsin solution 12.5 ng/µL in 50 mM NH_4_HCO_3_ (trypsin porcine, PROMEGA) for 1h on ice and incubated in 50 mM NH_4_HCO_3_ overnight at 37°C. The peptides were extracted twice with 50 µL of acetonitrile/water/formic acid-45/45/10-v/v/v followed by a final extraction with 50 µL of acetonitrile/formic acid (FA)-95/05-v/v. Peptides were dried in a speed-vac before nano LC-MS/MS analysis and then suspended in 0.1% HCOOH.

The sample was analyzed using an Ultimate 3000 nano-RSLC (Thermo Scientific, San Jose California) coupled on line with a Q Exactive HF mass spectrometer via a nano-electrospray ionization source (Thermo Scientific, San Jose California).

5 µL of peptide mixtures was loaded on a PepMap NEO C18 trap-column (300 µm ID x 5 mm, 5 µm, Thermo Fisher Scientific) for 3.0 minutes at 20 µL/min with 2% ACN, 0.05% TFA in H_2_O and then separated on a C18 Acclaim Pepmap100 nano-column, 50 cm x 75 mm i.d, 2 mm, 100 Å (Thermo Scientific) with a 40 minutes linear gradient from 8.8% to 50% buffer B (A: 0.1% FA in H_2_O, B: 0.1% FA in ACN), from 50 to 90% of B in 0.5 min, hold for 4 min and returned to the initial conditions in 0.5 min for 15 min. The total duration was set to 60 minutes at a flow rate of 300 nL/min. The oven temperature was kept constant at 40°C. Sample was analysed with TOP20 HCD method: MS data were acquired in a data dependent strategy selecting the fragmentation events based on the 20 most abundant precursor ions in the survey scan (375-1600 Th). The resolution of the survey scan was 60,000 at m/z 200 Th. The Ion Target Value for the survey scans in the Orbitrap and the MS^2^ mode were set to 3E6 and 1E5 respectively and the maximum injection time was set to 60 ms for both scan modes. Parameters for acquiring HCD MS/MS spectra were as follows; resolution 15,000 at m/z 200 Th, collision energy = 27 and an isolation width of 2 m/z. The precursors with unknown charge state or a charge state of 1 were excluded. Peptides selected for MS/MS acquisition were then placed on an exclusion list for 20 s using the dynamic exclusion mode to limit duplicate.

## Data Analysis

Proteins were identified by database searching using Sequest HT and MS Amanda with Proteome Discoverer 2.5 software (Thermo Scientific) against the RocS sequence and uniprot *S. pneumoniae* R6 database. Precursor mass tolerance was set at 10 ppm and fragment mass tolerance was set at 0.02 Da, and up to 2 missed cleavages were allowed. Oxidation (M), acetylation (Protein N-terminus), Phosphorylation (S, T, Y) were set as variable modification, and Carbamidomethylation (C) as fixed modification. Trypsin was selected as full and two miss-cleavage were allowed. False discovery rate for peptides and proteins was set at 1%. phosphopeptides identified in high confidence were checked manually.

### Electrophoretic mobility shift assay

Electrophoretic mobility shift assays (EMSA) were performed in 6% native polyacrylamide gel. The DNA fragment was amplified on the genome of *S. pneumoniae* R6 strain *via* PCR using primers CTCGGGACTTTTTTAAGTATGCCC and CCAACTGCTGGACGAGCTGC TAAG. Purified RocS-ΔAH (WT or point mutants) was mixed with 100 ng of DNA fragment (1000 :1 ratio) in a final volume of 20 μL. The binding buffer contained 25 mM Tris/HCl pH 7.5, 100 mM NaCl and 5% glycerol. The reaction was incubated for 30 minutes at room temperature. The polyacrylamide gels were pre-run at 4°C at 60 V in 0.5x Tris-Borate-EDTA buffer prior to the loading of the samples. The complexes were then resolved by migration at 4°C for 4 hours at 60V. DNA was visualized by bromure ethidium staining and imaging on a UV-transilluminator.

### Microscale thermophoresis (MST)

The affinity of RocS WT and mutants for DNA was measured using MST with a Monolith NT.115 Series instrument (Nano Temper Technologies). The DNA probe labeled with Cyanin 5 in 5’ (dnaC_Cy5) was reconstituted by annealing a 50bp forward primer (Cy5-GTCAACCAAGCTTACGAAGCGTCACAACCAGCTGATGAAATTATTGCTCA) labeled with Cyanine 5 at its 5’-extremity and its unlabeled complementary strand (TGAGCAA TAATTTCATCAGCTGGTTGTGACGCTTCGTAAGCTTGGTTGAC) at 95°C for 10 minutes with subsequent progressive cooling at room temperature. A mix of 10 nM DNA probe (1:1 v/v) with increasing concentrations of RocS (from 0.0061 to 200 μM) was loaded into standard Monolith NT.115 capillaries and MST was measured at 25°C in MST buffer (25 mM Tris/HCl pH 7.5, 100 mM NaCl, 5% glycerol) supplemented with 0.025% of Tween-20. The measurements were performed in triplicates and the analysis was carried out with the Monolith Software. The affinity constant (K_D_) was derived from the fluorescence change *’measured in function of RocS concentration. The data are represented here as the variation of the normalized fluorescence in function of RocS concentration (3 independent experiments). The normalized fluorescence corresponds to the ratio between fluorescence after thermophoresis and initial fluorescence values.

### Structures and Alphafold model analysis

We generated a model of RocS structure as a dimer bound to dsDNA with AlphaFold3 (Fig. 2C, 3A and 3B) (49). We searched for structural homologs of the wHTH domain (residues 1-55) in the protein bank database using the Dali server (http://ekhidna.biocenter.helsinki.fi/dali) (50). The structures of Par (8csh-A), RacA (5i44-B) and the AlphaFold model of RocS (ranked_0) were aligned based on their secondary structures using the MatchMaker tool from ChimeraX (https://www.rbvi.ucsf.edu/chimerax) (51).

### Sequence alignments

The 11 C-terminal amino acids (aa) of MinD, RocS and Par were aligned with MAFFT (52). The wHTH sequences from RocS (aa 1-55), Par (aa 1-51) and RacA (aa 1-50) were aligned according to their secondary structures (structures of Par (8csh-A) and RacA (5i44-B) and the AlphFold model of RocS (ranked_0) were entered as queries) using Promals3D (http://prodata.swmed.edu/promals3d) (53). The sequences of DUF 536-containing proteins were retrieved from InterPro and aligned with MAFFT (52). All alignments were visualized using Jalview (http://www.jalview.org) (54).

## Data availability

All strains, plasmids and row data generated in the course of this work are available from the author upon request.

## ACKNOWLEDGMENTS

We would like to thank Céline Freton for expert technical assistance for microscale thermophoresis experiments and members of the Grangeasse lab for fruitful discussions. We acknowledge the contribution of the Protein Science plateform of the “Structure Féderative de Recherche” Biosciences Gerland-Lyon-Sud (UMS344/US8), in particular Adeline Page and Frédéric Delolme for mass spectrometry analysis and Virginie Gueguen-Chaignon for technical advice. In addition, we would like to thank Linda Doubravová and Bohumil Kubeša who kindly shared that RocS was also found as phosphorylated in their phosphoproteomics data. This project has received funding from the European union’s Horizon H2020 research and innovation programme under the Marie Sklodowska-Curie grant agreement N°955626. It was also supported by grants from the Centre National de la Recherche Scientifique, Université Claude Bernard Lyon 1 and the Bettencourt Schueller Foundation.

